# Application of CRISPR/Cas9 nuclease in amphioxus genome editing

**DOI:** 10.1101/2020.07.29.227843

**Authors:** Liuru Su, Chenggang Shi, Xin Huang, Yiquan Wang, Guang Li

## Abstract

The cephalochordate amphioxus is a promising animal model for studying the origin of vertebrate due to its key phylogenetic position among chordates. Although transcription activator-like effector nucleases (TALENs) has been adopted in amphioxus genome editing, its labor intensive construction of the TALEN proteins limits its usage in many laboratories. We here report an application of the CRISPR/Cas9 system, a more amenable genome editing method, in this group of animals. Our data show that while co-injection of *Cas9* mRNAs and sgRNAs into amphioxus unfertilized eggs causes no detectable mutations at targeted loci, injections of *Cas9* mRNAs and sgRNAs at two cell stage, or of Cas9 protein and sgRNAs before fertilization can executes efficient disruptions of targeted genes. Among the nine tested sgRNAs (targeting five genes) co-injected with Cas9 protein, seven introduced mutations at efficiency ranging from 18.4% to 90% and four caused specific phenotypes in the injected embryos. We also demonstrate that monomerization of sgRNAs via thermal treatment or integration of sgRNAs into a longer backbone could increase mutation efficacies. Our study will not only promote application of genome editing method in amphioxus research, but also provide valuable experiences for other organisms in which the CRISPR/Cas9 system has not been successfully applied.

## 1. Introduction

The cephalochordate amphioxus represents a transitional group between invertebrates and vertebrates, showing most similarities with vertebrates in terms of either body plan, genome structure or embryogenesis among the living animals. But compared to vertebrates, amphioxus has not undergone extensive genome duplications (Putnam et al., 2008) and lacks paired eyes, bones, appendages, a sophisticated brain and most visceral organs. Because of these, amphioxus has be thought to be a promising model for studying the origin and evolution of vertebrate complexity (Bertrand and Escriva, 2011). To develop amphioxus as a model organism, several breakthroughs have been recently made, which include development of methods for manipulating amphioxus spawning under demands throughout a year (Li et al., 2013b; Li et al., 2012) and generating amphioxus mutants with the TALEN system (Li et al., 2014) and transgenes via the Tol2 transposon system (Shi et al., 2018). In the present study, we further report a highly efficient, more amenable genome editing method in this organism, using the clustered regularly interspaced short palindromic repeat/Cas9 (CRISPR/Cas9) system (Cong et al., 2013).

## 2. Materials and Methods

### 2.1. Amphioxus maintenance and image acquisition

Amphioxus *Branchiostoma floridae* were obtained from Dr. Jr-Kai Yu’s laboratory (Institute of Cellular and Organismic Biology, Academia Sinica, Taiwan, China), and maintained under previously described conditions (Li et al., 2012). Gametes were obtained using thermal-shock (20 °C to 26 °C) (Li et al., 2013a). Amphioxus transgenic line Tg *(mylz2: mCherry)* has been reported previously (Shi et al., 2018). Larvae were photographed under an IX71 microscope (Olympus, Japan), or a M165FC stereoscope (Leica, Germany).

### 2.2. In vitro Cas9 mRNA synthesis

pXT7-Cas9 (Chang et al., 2013) is a gift from China Zebrafish Resource Center, CZRC, Wuhan, China, and pCS2-nls-zCas9-nls (Jao et al., 2013) was obtained from Dr. Wenbiao Chen. They were linearized respectively using *XbaI* and *NotI,* purified with phenol-chloroform method, and used as templates to synthesize *Cas9* mRNA with the mMESSAGE mMACHINE T7 or SP6 Transcription Kits (Thermo Fisher Scientific). Synthesized mRNAs were treated with the Tango DNase supplemented by the kit, purified by phenol-chloroform method, dissolved in RNase-free water, and stored at −80 °C for use

### 2.3. sgRNA design and synthesis

All sgRNAs except *mCherry-sgRNA* were designed at web site (https://www.crisprscan.org/?page=sequence) according to roles used for zebrafish. Two different versions of gDNA backbones were tested in the study. Version 1 is a regularly used one (Hsu et al., 2013), while version 2 is modified from version 1 by including a 5 bp duplex extension and a T-to-C mutation at position 4 of the continuous sequence of thymines (Dang et al., 2015). Version 1 gDNAs (DNA template used to synthesize sgRNA) were generated as follows. For each target site, a pair of primers containing target sequence and overhang were synthesized, annealed and ligated between two *BsmBI* sites of pT7-gRNA vector obtained from Dr. Wenbiao Chen (Jao et al., 2013). Resultant plasmids were used to amplify version 1 gDNAs with primers: M13R, 5’-agcggataacaatttcacacagg-3’; pT7-sgRNA-R1, 5’-gatccgcaccgactcggtgccact-3’. Version 2 gDNAs were obtained with a more straight-forward way by annealing a specific oligonucleotide containing the T7 promoter, spacer (target sequence) and partial repeat sequence, with a universal oligonucleotide containing whole repeat sequence. Both version 1 and 2 gDNAs were purified by phenol-chloroform method before being used as templates to synthesize sgRNAs with the MEGAshortscript™ T7 Transcription Kit (Thermo Fisher Scientific). The synthesized sgRNAs were further treated with the Tango DNase, purified by phenol-chloroform method, resolved in RNase-free water and stored at −80 °C for use. Primers and oligonucleotides used were listed in Supplementary Table S1, and sequence information of sgRNA backbones was provided in Supplementary Table S2.

### 2.4. Injection solution preparation and microinjection

*Cas9* mRNA/sgRNA injection solution was prepared to 5 mg/ml fluorescein-labeled dextrans (10,000 MW, Thermo Fisher Scientific), 20% glycerol, 376.7 μg/μl *Cas9* mRNA, and 178.6 μg/μl sgRNA. Cas9 nuclease/sgRNA injection solution was prepared to contain 3 mg/ml fluorescein-labeled dextrans, 12.5% glycerol (not including the glycerol from Cas9 nuclease) and desired concentration of Cas9 protein and sgRNA. Three Cas9 nucleases, which were respectively purchased from New England BioLabs (20 μM, equal to around 3 μg/μl), Thermo Fisher Scientific (1 μg/μl) and TaKaRa (3 μg/μl) were tested in the study. sgRNA was used directly or pre-denatured as previously described (Dang et al., 2015). Before injection, the Cas9 nuclease/sgRNA injection solution was incubated at 37 °C for 5 min in a thermal cycler to facilitate Cas9/sgRNA RNP complex formation. Microinjection of unfertilized eggs was carried out as previously described (Liu et al., 2013), while blastomere injection at the two-cell stage was conducted as recently described (Zhu et al., 2020).

### 2.5. Mutant efficiency estimation

Genomic DNA of embryos (~20) at and after 32-cell stage were extracted using Animal Tissue Direct PCR kit (FOREGENE) and those (~200) of unfertilized eggs and 2-cell embryos were purified using DNeasy^®^ Blood & Tissue Kit (Qiagen). DNA fragments flanking target sites were amplified using primers listed in Supplementary Table S3. Mutations were detected using either T7EI cleavage assay or restriction enzyme digestion assay. For some experiments, mutations were also confirmed by DNA sequencing. For T7EI cleavage assay, 20-40 ng amplicons were melted and reannealed in a 10 μl reaction system as follows: 95 °C for 5 min, 95 °C to 20 °C ramping at 0.1 °C /s and holding at 20 °C, and then digested by 2 U of T7 endonuclease I (T7E1, New England BioLabs) at 37 °C for 15 min; and for restriction enzyme digestion assay, 20-40 ng amplicons were digested in a 10 μl reaction system with proper enzymes. The digested samples were then subjected to electrophoresis on a 2% agarose gel. Mutation efficiencies were either estimated by comparing band intensity between uncut and cut bands (for restriction enzyme digestion assay), or calculated using formula: % mutation = 100 × (1-(1-fraction cleaved)^1/2^) (Guschin et al., 2010). Quantification of band intensity was executed with software implemented in Tanon Gis system (Tanon).

### 2.6. Quantitative RT-PCR

Injection solutions containing 511 ng/μl mCherry-sgRNA and 302 ng/μl *Cas9* mRNA, or 511 ng/μl mCherry-sgRNA and 906 ng/μl TaKaRa Cas9 protein were used in this experiment. After injection, thirty eggs or embryos were collected at unfertilized egg, 2-cell, and early gastrula stages. Their total RNAs were then extracted using TRIzol reagent (Life Technologies) and reverse-transcribed to cDNA using Evo M-MLV One Step RT-PCR Kit (Accurate Biotechnology (Hunan) Co.). Real-time quantitative PCR (RT-qPCR) analysis was performed using TransStart Tip Green qPCR SuperMix kit (TransGen Co.) on a Stratagene Mx3000P real-time PCR system (Stratagene, La Jolla, CA, USA) under the conditions of 94 °C for 30 s, 40 cycles at 94 °C for 5 s, 58 °C (for *Cas9* and *Gapdh)* or 61 °C (for *Gapdh* and *sgRNA*) for 15 s, and 72 °C for 10 s. Expression levels of sgRNA and *Cas9* were normalized to that of *Gapdh* (glyceraldehyde-3-phosphate dehydrogenase), and the 2-ΔΔCt method was used to calculate the relative expression of the two genes examined. Three independent experiments were conducted and results from them were used to calculate the averages and standard errors. The primer sequences used are listed in Supplementary Table S4.

## 3. Results and Discussion

### 3.1. Injection of Cas9 mRNA and sgRNA at 2-cell stage, but not at unfertilized egg stage, induces targeted mutations in amphioxus

We first tried injecting amphioxus eggs with *Cas9* mRNA and sgRNA as it was described in other species. Two versions of *Cas9* mRNA from pXT7-hCas9 and pCS2-nls-zCas9-nls vectors which have been shown to be highly efficient in zebrafish (Chang et al., 2013; Jao et al., 2013), and a previously reported sgRNA targeting *mCherry* gene (hereafter, *mCherry-sgRNA)* (Fig. S1A) (Hashimoto and Takemoto, 2015) were tested. After mixing each of these *Cas9* mRNAs with the mCherry-sgRNA we injected them respectively into unfertilized eggs of *mylz2: mCherry* transgenic amphioxus (Shi et al., 2018). Unexpectedly, we found no detectable mutations at the target site for either injection by T7EI cleavage assay (Fig. 1A) and DNA sequencing.

**Figure 1.**
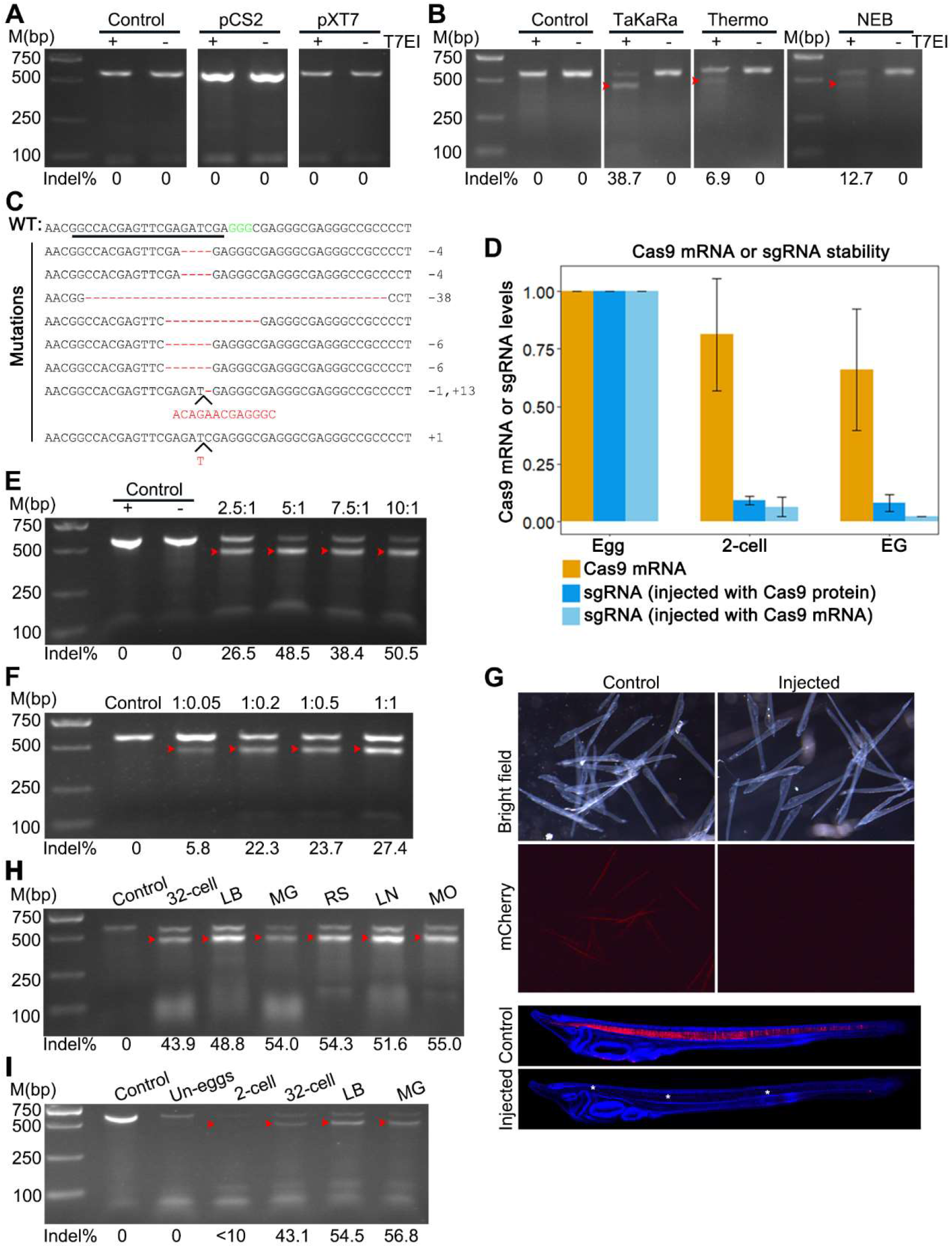
Detection of CRISPR/Cas9-induced mutation at *mCherry* locus in *mylz2-mCherry* transgenetic amphioxus. (A) T7EI cleavage assay showing no detectable mutation in uninjected embryos (control), or embryos injected with *mCherry*-sgRNA and *Cas9* mRNA transcribed from pXT7-Cas9 (pXT7) or pCS2-nls-zCas9-nls (pCS2) plasmids. – means no T7 endonuclease I was added and + means T7 endonuclease I was added. (B) T7EI cleavage assay showing no detectable mutation in uninjected embryos (control) and different levels of mutations in embryos respectively injected with mCherry-sgRNA and each of the three commercial Cas9 protein (TaKaRa, Thermo and NEB). – means no T7 endonuclease I was added and + means T7 endonuclease I was added. (C) Mutations detected by DNA sequencing in the injected embryos. The wild-type (WT) reference sequence is shown on the top. Sequence underlined is the target site and GGG (green) is the PAM sequence. Deletion is showed by dashed line and insertion is highlighted by inserted letters. Indels (+, insertion; -, deletion) are listed on the right of each allele. Red arrowheads in all above panels mark bands released by T7 endonuclease I digestion. (D) RT-qPCR analysis of *Cas9* mRNA and mCherry-sgRNA expression in embryos injected with Cas9 protein and mCherry-sgRNA, or *Cas9* mRNA and mCherry-sgRNA. (E) Mutation efficiencies in embryos injected with different molar ratio of TaKaRa Cas9 to *mCherry-sgRNA,* in which amount of Cas9 protein was kept constantly while that of *mCherry-sgRNA* was different. (F) Mutation efficiencies in embryos injected with different molar ratio of TaKaRa Cas9 to mCherry-sgRNA, in which amount of *mCherry-sgRNA* was kept constantly while that of Cas9 protein was different. (G) Observation of red fluorescent signal in 2-day transgenetic larvae. In contrast to uninjected (control) larvae which show specific fluorescent signal in the notochord and somites, the injected embryos show no detectable fluorescent signal. Blue fluorescent signal shows the amphioxus cell nuclei stained with DAPI. H-I: Mutation efficiencies in different stages of embryos injected with TaKaRa Cas9 and mCherry-sgRNA. Un-eggs, unfertilized eggs; LB, late blastula; MG, mid-gastrula; RS, rotation stage; LM, late neurula; MO, mouth-opening stage.

We speculated that this negative result might be caused by degradation of sgRNA in amphioxus embryos, since less efficacy led by crRNA degradation has been reported in the application of the Cpf1 system in zebrafish embryos (Moreno-Mateos et al., 2017). To test this, we monitored the *mCherry*-sgRNA level in three different stages of embryos by RT-qPCR. Consistent with our speculation, we observed that the *mCherry*-sgRNA level rapidly decreased after injection and only about 6 percent of them was retained at the 2-cell stage (about 50 minutes after injection). The level further reduced to around 2% at the early gastrula stage (Fig. 1D). To avoid the major decay of sgRNA occurred between unfertilized egg and 2-cell stages, we injected *Cas9* mRNAs with sgRNAs into one blastomere at 2-cell stage. Four sgRNAs were tested and three of them (except for the mCherry-sgRNA) induced mutations at their target sites with efficacies ranging from 21% to 33.3% (Fig. S1C). These results demonstrated that sgRNA decay occurred before 2-cell stage is a major cause for ineffectiveness of coinjection of *Cas9* mRNAs and sgRNA at the unfertilized egg stage.

### 3.2. Effective mutations caused by injecting Cas9/sgRNA ribonucleoprotein (RNP) complexes in amphioxus unfertilized eggs

Since injection of amphioxus embryos at 2-cell stage is much more difficult and often leads to more embryo deformation than injection of amphioxus unfertilized eggs, we further tried to inject Cas9/sgRNA ribonucleoprotein (RNP) complexes at the unfertilized egg stage. We speculate that the incorporation of sgRNA into Cas9 protein would prevent or at least slow down the sgRNA decay observed in above experiments. We tested three kinds of commercial Cas9 nucleases, which were respectively purchased from TaKaRa, Thermo and NEB companies, at the mCherry-sgRNA target site. We detected mutations in embryos injected with each of the three Cas9 nucleases using T7EI cleavage assay and DNA sequencing (Fig. 1B, Supplementary Figure S1B). Among them, the nuclease from the TaKaRa company shows the highest mutation efficacy (38.7% vs 12.7% and 6.9%) (Fig. 1B). The mutations induced by Cas9 protein from TaKaRa company include insertions and deletions of sizes ranging from 1 bp to 38 bp (Fig. 1C). Interestingly, we also detected a substantial decrease in the sgRNA level in this experiment (TaKaRa Cas9 was used) at the 2-cell stage (Fig. 1D). But different from injection of *Cas9* mRNA, no further degradation of the sgRNA was observed in injection of Cas9 protein after the 2-cell stage (Fig. 1D). We chose the Cas9 protein from TaKaRa company for the following experiments.

We next examined the effect of the ratio of Cas9 to sgRNA on the efficiency of genome editing. We found when keeping mCherry-sgRNA amount constant (200 ng/μl), a twofold increase of Cas9 protein from 500 ng/μl (Cas9: *mCherry-sgNRA* = 2.5: 1) to 1000 ng/μl (Cas9: mCherry-sgNRA = 5: 1) could induce a significantly higher mutation efficacy (26.5% vs 48.5%) (Fig. 1E). However, further increasing the Cas9 amount (7.5: 1 and 10:1) failed to induce more mutations (Fig. 1E). A similar case happened when we kept Cas9 protein amount constant (500 ng/μl) and sgRNA amount variable. A fourfold increase of sgRNA from 25 ng/μl (Cas9: mCherry-sgNRA = 1:0.05) to 100 ng/μl (Cas9: *mCherry*-sgNRA = 1: 0.2) could induce nearly a four-fold increase in mutation efficacy (5.8% vs 22.3%), and further increasing the sgRNA amount (1: 0.5 and 1: 1) was unable to cause more mutations (Fig. 1F). These results indicated that the best weight ratio of Cas9 to sgRNA in amphioxus genome editing is 5: 1. Interestingly, this weight ratio corresponds a 1: 1 molar ratio, consistent with the finding that Cas9 protein and sgRNA form RNP complex in a 1: 1 molar ratio manner. Moreover, the embryos injected with a 5: 1 ratio of Cas9 (1000 ng/μl) to mCherry-sgRNA (200 ng/μl) failed to emit red fluorescence at 3-gill slit stage, while in uninjected larvae, around 50% of them show red fluorescence in their somites and notochords (Fig. 1G). This result again demonstrates a high mutation efficacy for this combination of the Cas9 and *mCherry-sgRNA.*

We then determined the dynamics of mutation efficiency along the development of embryos injected with the Cas9 and mCherry-sgRNA RNP complexes. We found no detectable mutation in injected unfertilized eggs, and less than 10% mutation efficacy at two-cell stage (around 1 hour post fertilization, 1 hpf) (Fig. 1I). By 32-cell stage (around 2 hpf), around 43.9% of the target site has been mutagenized; at late blastula stage, the mutation efficacy reached the peak (around 50%) (Fig. 1H and I). Therefore, compared to TALENs (Li et al., 2014), CRISPR/Cas9 could induce mutations in amphioxus much faster.

### 3.3. The broad feasibility of the CRISPR/Cas9 system in amphioxus genome editing

To test whether the Cas9/sgRNA system is of broad spectrum in genome editing in amphioxus, we synthesized eight other sgRNAs targeting coding sequences of *Hedgehog (Hh)* (one), *Dkk1/2/4* (two), and *Hwa* (two), and *Nodal* (three) genes (Fig. 2A, Fig. S2A), and injected each of them with the TaKaRa Cas9 protein into amphioxus embryos. Among these genes, function of *Hh* gene has been clarified by TALEN method (Hu et al., 2017), role of *Nodal* has been addressed by chemical inhibitor (Li et al., 2017), and those of *Dkk1/2/4* and *Hwa* genes have never been studied. We detected mutations in embryos injected with *Hh*-sgRNA, *Dkk1/2/4*-sgRNA1, *Dkk1/2/4-* sgRNA2, *Hwa*-sgRNA1, *Nodal*-sgRNA1 and *Nodal*-sgRNA2 at efficacies ranging from 18.4% to 90% (Fig. 2B and C, Fig. S2C). By contrast, we detected no mutations for *Hwa*-sgRNA2 and *Nodal*-sgRNA3 by restriction enzyme digestion assay (Fig. 2C, Fig. S2C) and DNA sequencing (Fig. S2B). Moreover, we observed specific phenotypes in embryos injected with *Hh*-sgRNA, *Nodal*-sgRNA1 and *Nodal*-sgRNA2 (Fig. S2D and E). These phenotypes (left isomerism and curled tail for *Hh*-sgRNA and right isomerism for *Nodal*-sgRNAs) are similar to those reported in embryos deficient of *Hh* gene (Hu et al., 2017) or Nodal signaling activity (Li et al., 2017). We also injected *Nodal*-sgRNA1 and *Nodal*-sgRNA2 simultaneously, and found this caused more embryos of *Nodal*-deficient phenotype than single sgRNA injections (Fig. 2E) although the mutation efficacies at each site are similar (Fig. 2D).

**Fig. 2.**
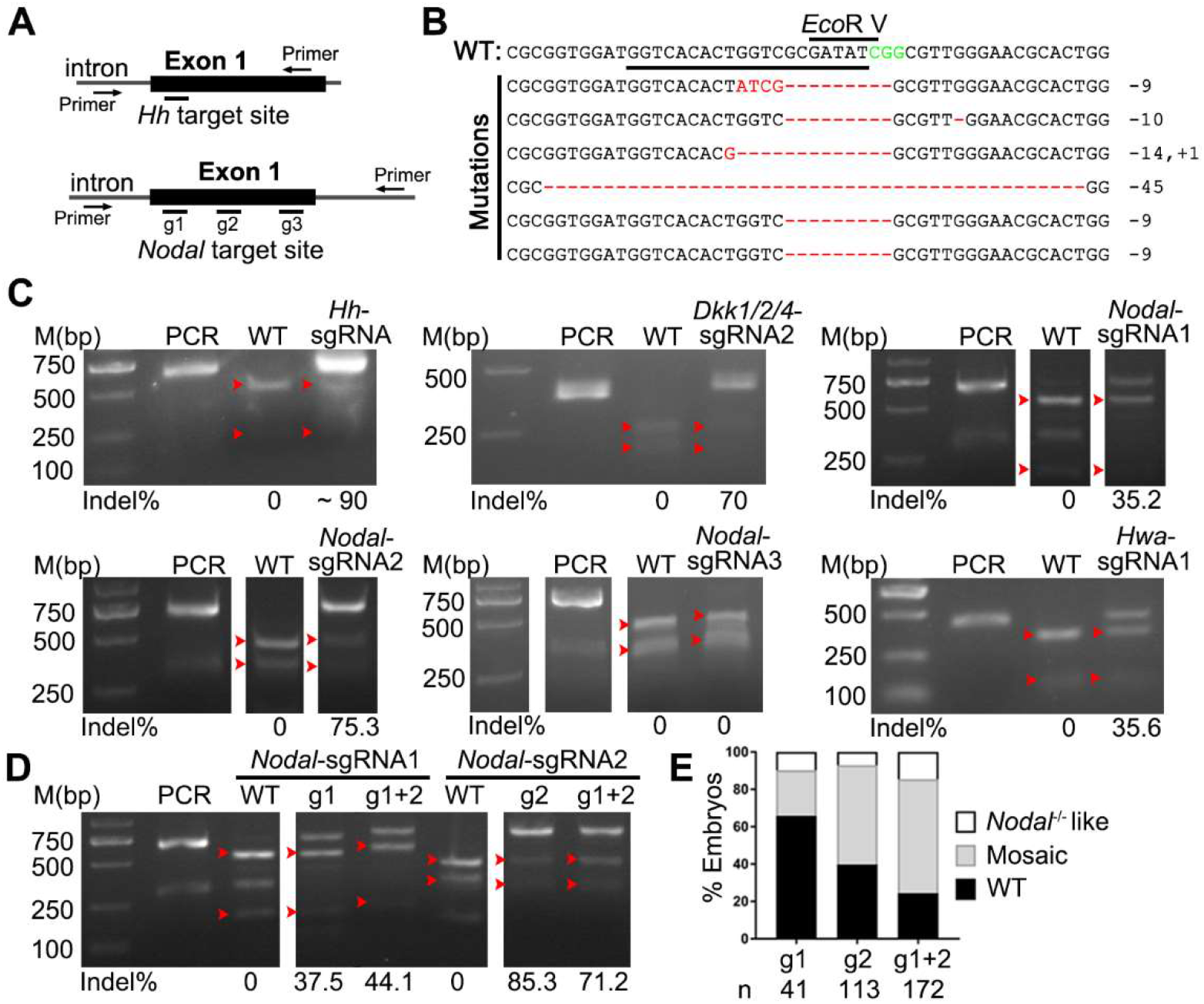
The broad feasibility of CRISPR/Cas9 system in genome editing of amphioxus. PCR, PCR products not treated with restriction enzyme; WT, digestion of PCR products from control (uninjected) embryos with restriction enzyme. Red arrowheads indicate bands released by restriction enzyme digestion. The induced mutation ratios (estimated as percentages of uncut PCR products) are labeled under each gel image. (A) Schematic showing the position of the target site and the primers used for PCR amplicons in the *Hh* and *Nodal* loci in amphioxus. (B) Sanger sequencing results showing mutations (red) mediated by Cas9/sgRNA in *Hh* allele. The wild-type (WT) reference sequence is shown on the top. Sequence underlined is the target site and CGG (green) is the PAM sequence. (C) Mutations detected by restriction enzyme digestion assay in the *Hh, Dkk1/2/4, Nodal,* and *Hwa* loci. (D) Comparison of mutation efficacies between single sgRNA and dual sgRNAs injections in the *Nodal* locus. (E) Phenotypic evaluation of embryos injected with single sgRNA or dual sgRNAs targeting the *Nodal* gene. Two kind of phenotypes are counted: embryos of *‘Nodal^-/-^-like* phenotype exhibit a left isomerism morphology resembling embryos lacking Nodal signaling, and embryos of ‘mosaic’ phenotype show left-right defects in positioning of some organs. Numbers (n) of embryos evaluated is shown for each condition.

### 3.4. Improvement of CRISPR/Cas9-mediated mutation efficiency in amphioxus

The *in vitro* synthesized sgRNAs exist in two bands in agarose gel (Fig. 3A), which are probably resulted from dimerization or secondary structure. We refer the short one (~100bp) to monomer and the long one (~200bp) to dimer for simplicity according to a previous report (Dang et al., 2015). Denaturation of sgRNA by a heating and quick cooling step could change the dimer to monomer and increase mutation efficacy (Dang et al., 2015). To test this in amphioxus, we denatured the synthesized sgRNAs before injection and compared their efficacies with the un-denatured ones. The treatment could efficiently reduce the proportion of dimers in all three tested sgRNAs (*mCherry*-sgRNA, *Dkk1/2/4*-sgRNA2 and *Hwa*-sgRNA1) (Fig. 3A), and induce more mutations at their target sites than the un-denatured ones (Fig. 3A). Especially, at the site targeted by Hwa-sgRNA1, the mutation efficacy was dramatically increased from 21.7% to 69.7%. Interestingly, this high level of efficacy increment correlates well with an highly efficient turnover of Hwa-sgRNA1 from dimer to monomer by the treatment (from less than 5% to around 40%) (Fig. 3A). This result demonstrates that a pre-denaturation of sgRNAs before mixing them with Cas9 protein could increase mutation efficacies in amphioxus, which is especially true for sgRNAs which tends to form dimers.

**Fig. 3.**
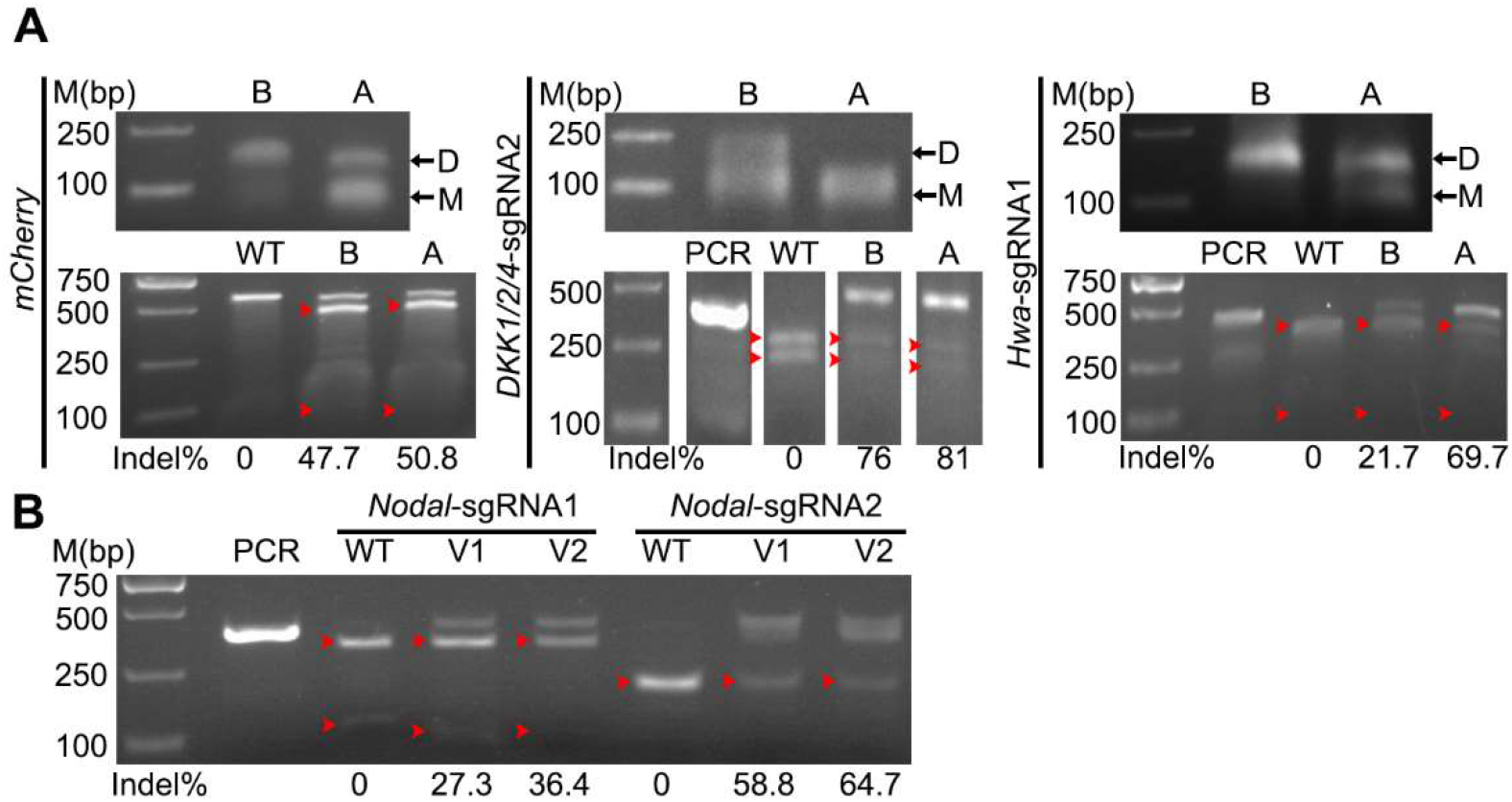
Improvement of mutation efficiency of Cas9/sgRNA in amphioxus. The induced mutation ratios shown under each gel image are estimated by either T7 endonuclease assay (for *mCherry-sgRNA)* or restriction endonuclease assay (for sgRNAs targeting the *Dkk1/2/4, Nodal* and *Hwa* genes). Red arrowhead indicates bands released by endonuclease digestions. PCR, PCR products not treated with restriction enzyme; WT, digestion of PCR products from control (uninjected) embryos with restriction enzyme. (A) Monomerization of sgRNAs (top) and its effect on mutation efficiency (bottom). sgRNA was monomerized by a denaturing step. B, before denaturation; A, after denaturation; D, dimer, M, monomer. (B) Comparison of mutation efficacies between version 1 (V1) and version 2 (V2) sgRNAs targeting the *Nodal* gene.

Although the sgRNA backbone used in above experiments is already an optimized one (Hsu et al., 2013) and has been widely used in many previous studies, a recent study showed that inclusion of a 5 bp duplex extension and a T-to-C mutation at position 4 of the continuous sequence of thymines in above backbone could further improve genome editing efficiency in cells (Dang et al., 2015). To test whether this is a similar case in amphioxus, we synthesized the two *Nodal*-sgRNAs of the newly updated backbone (named as version 2), and compared their efficacies to the original ones (named as version 1). The result showed that the two version 2 sgRNAs could both induce slightly more mutations than the original ones (36.4% vs 27.3% for *Nodal*-sgRNA1, 64.7% vs 58.8% for *Nodal*-sgRNA2) (Fig. 3B).

## Conclusions

In summary, we have successfully introduced the CRISPR/Cas9 system into amphioxus and demonstrated that it is able to induce targeted mutations highly efficient in this emerging model organism. Considering the simplicity of the CRISPR/Cas9 system, our work will certainly motivate more researchers to use genome editing method to study gene function in amphioxus. In addition, we found that nearly half of the tested sgRNAs (4/9) gave specific phenotypes among the injected embryos. This indicates a great potential for using it to knockdown gene function in amphioxus embryos. Moreover, our work also opens possibility for exploring other CRISPR/Cas9-based technologies, such as epigenetic modulation, inducible regulations, and base editing (Guo et al., 2018), in amphioxus.

## SUPPLEMENTARY DATA

Supplementary data to this article can be found online.

## COMPETING INTERESTS

The authors declare that they have no competing interests.

## AUTHOR CONTRIBUTIONS

G.L. and Y.W. designed and supervised the study. L.S., C.S., X.H., and G.L. performed the experiments and data analyses; L.S. and G.L. wrote the manuscript. All authors read and approved the final version of the manuscript.

## Funding

This work was supported by grants from the National Natural Science Foundation of China (No. 31872186, 31672246 and 31471986).

## Notes

### Competing Interest Statement

The authors have declared no competing interest.

