## Supplementary Figures and Tables for "Application of CRISPR/Cas9 nuclease in amphioxus genome editing"

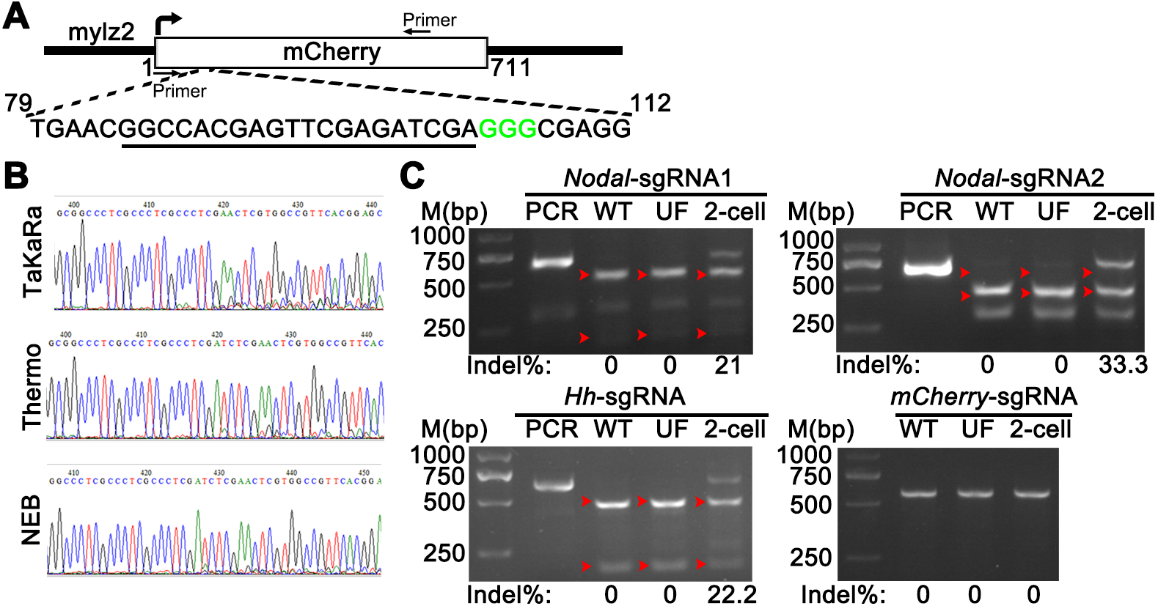

**Supplementary Figure S1 Cas9/sgRNA ribonucleoprotein complex induces site-specific mutations at *mCherry* locus in *mylz2-mCherry* transgenetic amphioxus**.

(A) Schematic showing the position of the target site and the primers used for PCR amplication in the *mCherry* gene. Sequence underlined is the target site and CGG (green) is the PAM sequence. (B) Sanger sequencing of *mCherry* locus from embryos injected with *mCherry*-sgRNA and Cas9 nuclease from three companies. (C) Enzyme digestion showing mutation induced by injecting Cas9 mRNA and corresponding sgRNAs at 2-cell stage.

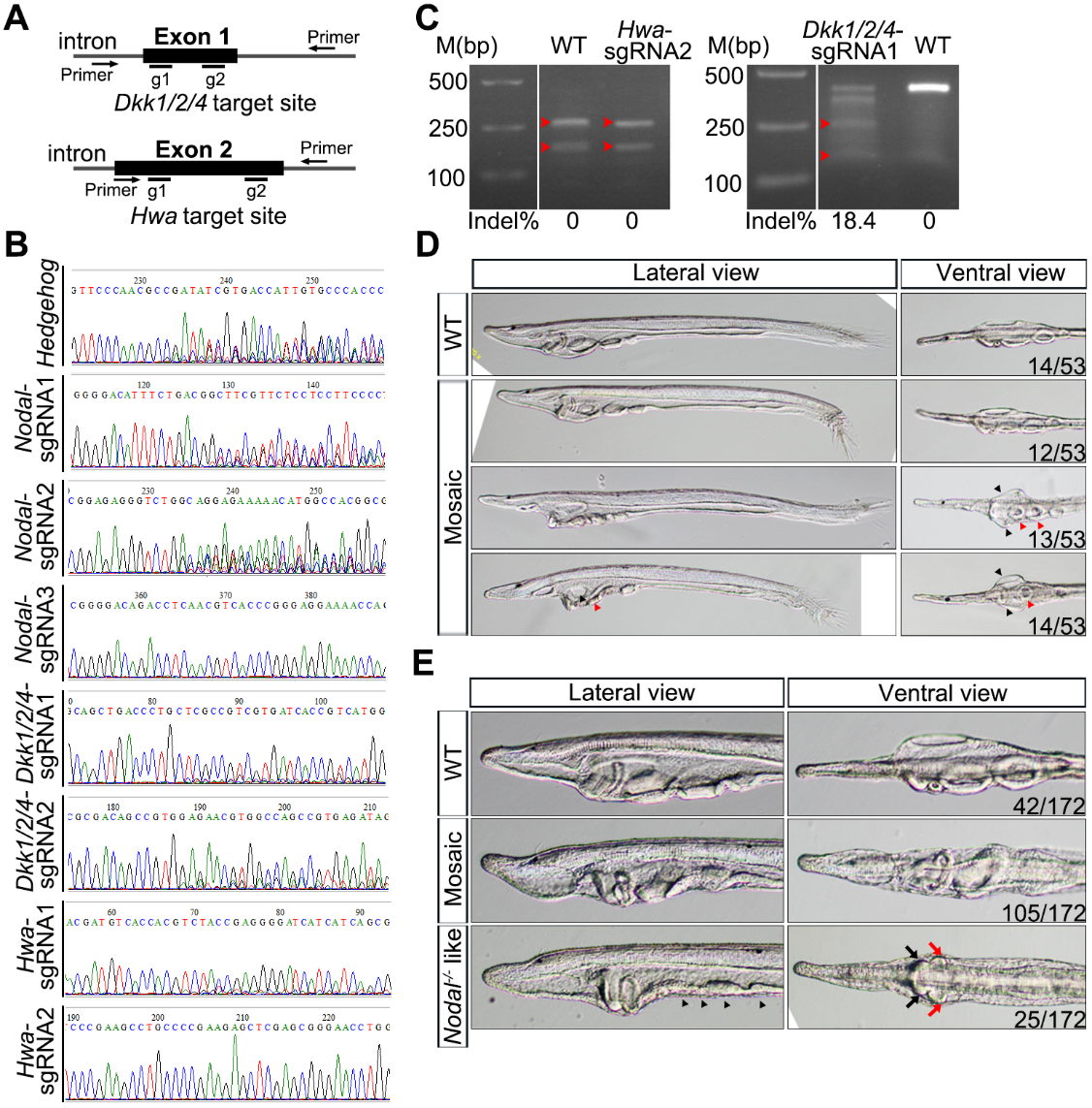

**Supplementary Figure S2 The broad feasibility of CRISPR/Cas9 system in genome editing of amphioxus**.

(A) Schematic showing the target sites and the primers used for PCR amplicons in the *Dkk1/2/4* and *Hwa* loci in amphioxus. (B) Sanger sequencing of amplicons of embryos injected with Cas9 nuclease and corresponding sgRNAs. (C) Mutation detection in embryos injected with Cas9 nuclease and *Hwa*-sgRNA2 (by restriction enzyme digestion assay) or *Dkk1/2/4*-sgRNA1 (by T7EI cleavage assay). Red arrowheads indicate bands released by endonuclease digestion. The induced mutation ratios are labeled under each gel image. (D) Representative phenotypes observed in 2-day larvae injected with Cas9/*Hh*-sgRNA RNP complexes. Among the 53 larvae examined, 14 were normal (WT) and 39 showed different level of similarities to *Hh*^-/-^ mutants (mosaic); out of the 39 ‘mosaic’ larvae, 12 developed a twisted tail (top), 13 left isomerism morphology (middle), and 14 showed both of the former phenotypes (bottom). It should be noted that none of these larvae lacked mouth like *Hh*^-/-^ mutants. Black arrowheads indicate the two symmetric mouths and red arrowheads indicate the ventralized gill slits that normally appear in the right side. (E) Representative phenotypes observed in 2-day larvae injected with Cas9 and mixture of *Nodal*-sgRNA1 and *Nodal*-sgRNA2 (version 1). Only the anterior portion of the larvae is shown, because no defects was observed in their posterior ends. Black arrowheads demarcate the ventralized and improperly developed gill slits, black arrows indicate the symmetric endostyles, and red arrows indicates the symmetric club-shaped gland.

**Table S1 Primers for construction of sgRNA vectors (V1 or V2)**

| Targeting  sites | Oligos (5’ to 3’) |
| --- | --- |
| *mCherry* | Forward: taggCCACGAGTTCGAGATCGA  Reverse: aaacTCGATCTCGAACTCGTGG |
| *Hedgehog* | Forward: taggTCACACTGGTCGCGATAT  Reverse: aaacATATCGCGACCAGTGTGA |
| *Dkk1/2/4*-sgRNA1 | Forward: taggTGATCACGACGGCGAGCA  Reverse: aaacTGCTCGCCGTCGTGATCA |
| *Dkk1/2/4*-sgRNA2 | Forward: taggCGACAGCCGTGGAGAACG  Reverse: aaacCGTTCTCCACGGCTGTCG |
| *Nodal*-sgRNA1 | Forward: taggGGACATTTCTGACGGCTT  Reverse: aaacAAGCCGTCAGAAATGTCC |
| *Nodal*-sgRNA2 | Forward: taggCGGAGAGGGTCTGACGCT  Reverse: aaacAGCGTCAGACCCTCTCCG |
| *Nodal*-sgRNA3 | Forward: taggACAGACCTCAACGTCACC  Reverse: aaacGGTGACGTTGAGGTCTGT |
| *Hwa*-sgRNA1 | Forward: taggTGTCACCACGTCTACCGA  Reverse: aaacTCGGTAGACGTGGTGACA |
| *Hwa*-sgRNA2 | Forward: taggTTCCCGCTCGAGCTCTTC  Reverse: aaacGAAGAGCTCGAGCGGGAA |
| pT7-gRNA-F1 | GGAATTAATACGACTCACTA |
| Dr-gRNA-R1 | AAAGCACCGACTCGGTGCCAC |
| universal primer | AAAGCACCGACTCGGTGCCACTTTTTCAAGTTGATAACGGACTAGCCTTATTTAAACTTGC  TATGCTGTTTCCAGCATAGCTCTTAAAC |
| BfNodal-gRNA1 | GGAATTAATACGACTCACTATAGGGGACATTTCTGACGGCTTGTTTAAGAGCTATGCTGG |
| BfNodal-gRNA2 | GGAATTAATACGACTCACTATAGGCGGAGAGGGTCTGACGCTGTTTAAGAGCTATGCTGG |
| mCherry-gRNA1 | GGAATTAATACGACTCACTATAGGCCACGAGTTCGAGATCGAGTTTAAGAGCTATGCTGG |

**Note**: red letters denote the target sequences

**Table S2 The sequences of different sgRNA structure used in this study**

| V1 | (N)_20_GTTTAGAGCTAGAAATAGCAAGTTAAAATAAGGCTAGTCCGTTATCAACTTGAAAAAGTGGCACCGAGTCGGTGCGGATC |
| --- | --- |
| V2 | (N)_20_GTTTAAGAGCTATGCTGGAAACAGCATAGCAAGTTTAAATAAGGCTAGTCCGTTATCAACTTGAAAAAGTGGCACCGAGTCGGTGCTTT |

**Table S3** **Primers used for PCR-based genotyping**

| Primers | Forward primer (5’ to 3’) | Reverse primer (5’ to 3’) |
| --- | --- | --- |
| *mCherry* | cggccgTGGTGAGCAAGGGCGAGGAG | TCTTGACCTCAGCGTCGTAGTG |
| *Hedgehog* | GCCGCTTTCTGCCTTGCTTCA | GCGAGTAATCCGTCCGTTGA |
| *Dkk1/2/4* | AGGATCTTCCCGAAACTCGC | AAGACGTACACCGGAAAGCC |
| *Nodal*-1 | CCGCCTTGCCTTTGTCTTTC | CACTGGTAGAAACGTCGTCCA |
| *Nodal*-2 | GGCAGGCCGAGACCAACACC | AGTGCGAAACTCCTGACGGTGTCG |
| *Hwa* | GAACAGGTGACTCAAACCCAAAG | TCTGGTCTCTGTGAAATGGCTTA |

**Note:** The lowercase in *mCherry* forward primer means the *Eag*I recognition sequence. The *Nodal*-2 Primers were only used in Fig. S4B for the reason that the *Nodal*-1 Primers were nonspecific at that experiment.

**Table S4 Primers used for qPCR**

| qPCR Primers | Forward primer (5’ to 3’) | Reverse primer (5’ to 3’) |
| --- | --- | --- |
| mCherry-sgRNA-RT | CACGAGTTCGAGATCGAGTTTT | CCGACTCGGTGCCACTTTTT |
| (WH)pXT7-hCas9-RT | AGGAGGACATCCAGAAAGCAC | TTCTCGGGCTTATGCCTTCC |
| Gapdh | GGTGGAAAGGTCCTGCTCTC | CTGGATGAAAGGGTCGTTAATGG |
